# Towards a system-level causative knowledge of pollinator communities

**DOI:** 10.1101/2021.09.23.461517

**Authors:** Serguei Saavedra, Ignasi Bartomeus, Oscar Godoy, Rudolf P. Rohr, Penguan Zu

## Abstract

Pollination plays a central role both in the maintenance of biodiversity and in crop production. However, habitat loss, pesticides, invasive species, and larger environmental fluctuations are contributing to a dramatic decline of numerous pollinators world-wide. This has increased the need for interventions to protect the composition, functioning, and dynamics of pollinator communities. Yet, how to make these interventions successful at the system level remains extremely challenging due to the complex nature of species interactions and the various unknown or unmeasured confounding ecological factors. Here, we propose that this knowledge can be derived by following a probabilistic causal analysis of pollinator communities. This analysis implies the inference of interventional expectations from the integration of observational and synthetic data. We propose that such synthetic data can be generated using theoretical models that can enable the tractability and scalability of unseen confounding ecological factors affecting the behavior of pollinator communities. We discuss a road map for how this probabilistic causal analysis can be accomplished to increase our system-level causative knowledge of natural communities.

## Introduction

Pollinators comprise a highly diverse group of species including bees, flies, butterflies, beetles, and some vertebrates [1]. They all have in common a shared interest in visiting flowers to extract resources, collectively and indirectly mediating the reproduction of most of the worldwide plant species [2] and maximizing crop production for 75% of cultivated crops [3]. Hence, pollination is now recognized not only as a key ecosystem function, but also as a key ecosystem service contributing to human food security. However, human induced rapid environmental change has been threatening most of these pollinators [4]. On the one hand, habitat destruction and modification is reducing the populations of many pollinator species, often leading to local extirpation. On the other hand, some other species can thrive in human modified ecosystems, but those often face extra pressures such as pesticide exposure, exotic species, or pathogens. In top of that, climate change is altering species’ physiological responses, distribution, and activity periods [5]. Overall, we are assisting to a rapid restructuring of pollinator communities world-wide, where their relative abundance, composition, and ecological interactions are being modified with hard to predict consequences for their health.

These human pressures on pollinator communities have increased the need for human interventions to protect the composition, functioning, and stability of pollinators and their interactions [6]. These interventions include from well established practices such as habitat protection, to more complex actions such as the addition or removal of particular species and their interactions [7]. For example, planting field margins [8] or adding managed pollinators [9] have become, respectively, popular restoration practices in agricultural systems to increase resources for pollinators or supplement crop pollination. However, these practices often ignore side effects, such as the effects of changes in micro-climate conditions or pathogen prevalence on pollinator health. For instance, a recent study has shown that bumblebees’ occupancy patterns in Europe and North America are sensitive to temperature [10]. Similarly, it has been shown how managed pollinator densities not only increases competition among pollinators [11], but also increases parasite loads [12], which can spillover to other species [13]. Yet, as of today, we lack a community-wide framework to guide interventions beyond single species. Indeed, it has been shown that even small local interventions (i.e., at the species level) can have heterogeneous and arbitrary cascading effects across entire communities [14]. This has emphasized the dire need to establishing a system-level causative knowledge of pollinator communities.

To address the challenge above, ideally, we need to establish well-defined experiments eliminating all sources of bias (e.g., using randomized controlled trials) and test the effectiveness of a given intervention [15]. However, those sources of bias become extremely difficult to identify and measure in changing natural ecological communities conformed by several co-occurring and interacting species [16]. Moreover, many of these interventions may not be ethical (e.g., species removal) or feasible to perform because pollinators move freely and are difficult to track. This implies that it is instead necessary to obtain interventional knowledge from observational data (e.g., field studies or partially controlled studies) using causal-inference analysis [17]. These observational data (that record for example the observed presence/absence of pollinators) differ from fully controlled studies (that remove or add pollinators) in the sense that observational variables are the result of what is perceived and not of what is intervened by the investigator. Importantly, these observational data are typically confounded by unknown factors (also known as noise, context, or environmental conditions), such as biotic and abiotic variables, making difficult to differentiate between spurious and actual cause-effect relationships. To circumvent this problem, we propose that interventional knowledge can be inferred from the integration of observational and synthetic data. These synthetic data can be generated using theoretical models that can enable the tractability (operationalization and reproducibility) and scalability (generalization across dimensions) of unseen confounding factors acting at the community level. This framework can provide a probabilistic knowledge of how likely is a given cause to generate a target effect within a pollinator community (i.e., focusing on the probability of causes instead of effects). In the reminder, we discuss a road map for how this probabilistic causal analysis can be accomplished and illustrate it with a case study.

## Observational data: known factors

Given the lack of systematically controlled experiments, observational data from field studies or quasi-controlled experiments (where few factors may be controlled) can provide the raw material to understand the behavior (e.g., composition, dynamics) of a community. This behavior comes in the form of a joint probability distribution P_**V**_ over a set of relevant variables **V**. For example, studies may record any aspect of community composition as a function of a set of semi-controlled variables such as the presence (or density) of specific pollinators [18], their floral resources including both the identity of interacting plant species [19] and plant chemical composition [20–22], top down regulators including pathogen [23] and predators [24], as well as several environmental variables such as temperature [25, 26] or pesticide exposure [27, 28]. These observational studies can be either for a specific period of time (across different locations) or measure pollinator communities repeatedly over time in order to capture a wider range of temporal conditions affecting pollinators’ population trajectories, which often follow non-linear dynamics [29, 30].

While observational data are designed to track potential mechanisms affecting pollinator communities, they cannot establish cause-effect relationships by themselves, only associations [15, 17]. That is, following Reichenbach’s principle [31], if two variables (*X*, *Y*) are statistically related, then there exists a third variable or set of variables (*Z*) that causally influenced both (known as confounding effect: *X* ← *Z* → *Y*). In some situations, *Z* coincides with either *X* or *Y* (i.e., *Z* = *X* or *Z* = *Y*), establishing a causal link between *X* and *Y* (i.e., *X* → *Y* or *Y* → *X*). But without knowledge of *Z* (or when this unknown effect cannot be blocked from the analysis), we cannot safely conclude cause-effect relationships. Thus, conditional distributions (e.g., P_*Y*|*X*_) derived from observational data can coincide with causal mechanisms (e.g., *X* → *Y*), but not necessarily. Similarly, two variables (*X*, *Y*) may be statistically related if both are the common (confounding) causes of a given effect *Z* (i.e., *X* → *Z* ← *Y*: known as collider in the causal-inference literature [15]) upon which the data is selected (known as selection bias). This problem typically arises when data is filtered or conditioned by *Z* and 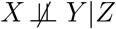, but 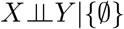 (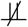 and 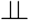 denote dependence and independence, respectively). Moreover, in a multivariate system, the sources of bias can be originated from direct and indirect common causes and effects. These properties make extremely problematic the interpretation of relationships derived from multivariate regression and meta-analysis that do not have a causal hypothesis [32].

For example, let us assume that pollinator abundance is caused by flower abundance, temperature, and some unknown factors. Similarly, let us assume that flower abundance is caused by water availability, temperature, and a subset of the same unknown factors. Then, in a multivariate regression model that includes all factors (except for the unknown) as potential explanations of pollinator abundance, it is likely that water availability will have a strong explanatory effect over pollinator abundance (even though we are conditioning over flower abundance). This happens for the reason that flower abundance introduces a selection bias (collider) between water and the unknown factors, which then gets propagated to pollinator abundance following the cause-effect relationships. Note that flower abundance cannot be eliminated from the regression model either, because it is needed to partially block the path between water availability and pollinator abundance. This type of examples also illustrates that prediction is different from explanation [33]. Therefore, to infer cause-effect relationships in this example, it is needed to have more information about the underlying causal story and the corresponding unknown confounding factors. In the next sections, we will discuss how to use synthetic data derived from theoretical models to account for confounding unobserved variables, and then how to generate interventional distributions (knowledge) from observational and synthetic data.

## Synthetic data: unknown factors

The role of theoretical models has been understood as a formal platform to establish logico-mathematical postulates (formal statements) about how the real-world possibly behaves and to obtain data that can be difficult to generate empirically [34–36]. These postulates are, of course, tautological as they are analytically (or algorithmically) derived from a set of primary principles. It is only possible to falsify these postulates based on their biological interpretation. Thus, the value of theoretical models is to provide hypotheses, predictions, generalization, and systematic links between model parameters (the interpretable factors/context) and the behavior of a system, which can then be revised based on empirical information. The interpretation of theoretical models (model parameters) can range from highly mechanistic to highly phenomenological depending on the level of resolution under investigation [37]. For example, mechanistic interpretations are based on detailed descriptions of ecological processes, such as metabolic rates, nutrients uptake, mobility patterns, predation processes, and behavioral patterns, among others [38, 39]. In turn, phenomenological interpretations are based on summary outcomes that are expressed in terms of model parameters without establishing any specific statement about how exactly these outcomes come to existence (e.g., intrinsic growth rates, species interactions, and death rates, among others). In general, there is no one better model than another (unless there is knowledge about the actual processes and there is capacity to obtain the initial conditions), it all depends on the research question and system under investigation.

Regarding pollinator communities (and ecological communities in general), there are two important properties that need to be considered if one aims to study theoretically and systematically the factors under which several interacting species can coexist [40]: tractablity and scalability. We define tractability as the property of a theoretical model to have all its potential solutions fully operationalized, defined, measured, and reproduced over relatively short periods of time (i.e., polynomial time), enabling a systematic understanding between solutions and parameter values. For example, the Londsdorf [41] model uses only land use parameters to directly explain pollinator densities following a simple equation. Instead, complex models characterized by higher-order polynomials are limited by their intractability (e.g., optimal foraging models [40, 42, 43]). In fact, it has already been proved that it is impossible to write analytically (a closed-form algebraic solution) a polynomial system with degree five or higher with arbitrary coefficients (unknown values) [44]. Note that a simple 3-species system (e.g., two pollinators and one plant) with Type II functional responses (i.e., a non-linear response such as those observed in density-dependent processes arising from competition for floral resources or pathogen spillover) can already form a polynomial of degree eight [45]. This intractability of complex models implies that if the majority of their parameter values are not known a priori (reducing the system to a polynomial of degree four or lower), these models can only be used numerically (simulations). Then, the problem that arises is that it becomes computationally impossible to differentiate the role played by each parameter (e.g., interactions, environmental conditions) in the solutions of the system [40]. While studies have attempted to tackle this complexity by using statistical methods such as Akaike Information Criterion [46], the number of solutions of a polynomial system does not necessarily depend on the number of parameters but on the polynomial degree [45]. Hence, it is not just the lack of data that limits the use of complex models, as it can be perceived [47], it is their intractability, especially in high-dimensional systems [40].

In turn, we define scalability as the property of a model to establish clear and invariant rules across dimensions, enabling extensions from simple to complex natural communities. For example, the Lonsdorf model [41] is designed to track central place foragers (e.g., bees), where a key piece of the model is the foraging range from a central point in the landscape; but it is not scalable to wanderers (e.g., flies and butterflies), which move freely over the landscape tracking resources. Similarly, it has been demonstrated that insights derived from classic work on coexistence using 2-species Lotka-Volterra models cannot be directly extrapolated to higher dimensions [48]. Therefore, simple phenomenological or simple mechanistic models can be understood as the simplification (reduction of polynomial degree and free parameters) of complex models to enhance the tractability and scalability of a system. However, it is central to fully understand how they should be used.

For instance, generic phenomenological models can be written in the form **Ṅ** = **N***f*(**N**, **U**), where **Ṅ** represents the time derivative of species density, and *f* is a given function describing the relationship among endogenous **N** variables and contextual parameters **U** [36]. Note that having the vector **N** in front of the function *f* guarantees the impossibility of negative densities (or species revival without immigration). A classic phenomenological model that follows this formalism is the linear Lotka-Volterra (LV) model [49, 50]: **Ṅ** = **N**(**r** + **AN**), where **r** typically represents species intrinsic growth rates and **A** is the so-called interaction matrix (summarizing the positive or negative per capita effect of one species upon individuals of another species). While the linear LV model can be derived from first principles, such as energy conservation or thermodynamic limits, it can be phenomenological interpreted as the first-order approximation (derived from the Taylor expansion) of the unknown function *f* [35]. This can then make the elements of the linear LV model to be interpreted as endogenous variables **N**, a set of time-invariant interaction parameters summarized in **A**, and contextual parameters **r**. This interpretation allows both the tractability and scalability of a multispecies community. That is, the analytical solution is **N**\* = −**A**^−1^**r** (setting **Ṅ** = 0), making possible the one-to-one mapping between **N**\* and **r** [51]. This means that the constraints imposed by **A** on contextual factors **r** to generate a given endogenous behavior **N**\* can be systematically analyzed regardless of the number of species in the system.

Importantly, tractable and scalable models become good candidates towards increasing our systemlevel causative understanding of pollinator communities. Indeed, by conceptualizing the function *f* above as an approximation to a structural causal model [15, 17] (i.e., *X* = *f_X_*(**V_X_**, **U_X_**), where *f_X_* is a time-invariant function defining the cause-effect relationships of *X*, **V_X_** is the set of causes of *X*, and **U_X_** is the random noise/context affecting *X* defined by P**_U_X__**), it is possible to obtain theoretical probability distributions of unknown factors **U** (e.g., **r** in the LV model) compatible with a given behavior of **N**\* as dictated by a set of invariant rules (e.g., **A** in the LV model).

For example, in the linear LV model, by assuming that 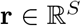 (where *S* is the dimension of the system) is a priori randomly and uniformly distributed (conforming with ergodicity and independence from initial conditions [52]), it is possible to calculate analytically the range of feasible unknown conditions (i.e., **U** ⊆ **r** and P_**U**_) leading to a given set of species (i.e., *I* ⊆ *R*, where *R* is the set of species within a community) with positive densities at equilibrium 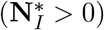 [53, 54]. Moreover, we can calculate the expected number of species with positive densities at equilibrium *E*[**N**\* > 0] (or the probability of persistence of each single species within a community) [52]. Note that if **A** is also derived from a probability distribution (i.e., P_**A**_), the range of feasible unknown conditions remains characterized by P_**U**_. Importantly, extracting these conditions requires the inference (empirical parameterization) of invariant rules (e.g., **A**). While challenging, it has been shown that this properties can be approximated with commonly available data, such as species abundances or presence/absence data [14, 55–58]. We provide a case study in the last section.

## Probability of causes

While observational data per se are not enough to obtain a causative knowledge about pollinator communities, they can be translated into interventional distributions using causal-inference techniques [15, 17]. Recently, promising causal-inference methods have been developed, such as inverse modelling approaches [59, 60] or empirical dynamical modeling [61], but these methods require large amounts of data which for several reasons can be difficult to obtain. To partially circumvent this problem, we propose that probabilistic causal-inference approaches [15] used in economics, social science, and medicine can be good candidates for inferring interventional distributions (i.e., how likely is a given cause to generate a target effect) in pollinator communities.

First and foremost, probabilistic causal inference requires a causal graph involving the set of relevant variables (nodes) **V** (e.g., **V** = {*X*, *Y*}, *X* → *Y*) upon which to test causal relationships (edges) [15]. These graphs serve as a guideline (testable hypothesis) to understand the potential paths linking causes and effects, which are necessary to study in order to eliminate spurious associations (due to confounding and selection bias). In general, causal graphs should be drawn based on expert knowledge or intuition about how the world works, and should not be drawn based on the observed correlations on data (otherwise, it will be circular). These graphs act as a hypothetical causal story, which can be followed after identifying and corroborating its testable implications expressed as unconditional and conditional independencies between variables (in causal-inference analysis, this is called d-separation of variables [15]). For instance, a lack of correlation between two variables in any context does not immediately invalidate a potential direct causal link (since we cannot be sure of having sampled all potential values within the sample space); however, a lack of correlation in all contexts after conditioning by a potential confounder (i.e., 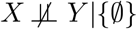, but 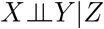) does support the hypothesized causal graph *X* ← *Z* → *Y* (i.e., no direct causal effect between *X* and *Y*). Remember that a correlation between two variables is not enough evidence to support a potential causal link. Thus, causal graphs inform about both the likely dependencies and established independencies between variables. If the data do not corroborate the causal graph, then a new causal story must be drawn and tested.

Causal graphs are nonparametric by construction since they do not depend on the specific form of causal relationships, they only specify the (lack of) existence of a causal relationship between variables. While most of the standard work on probabilistic causal inference has been developed for directed acyclic graphs (no mutual causality or feedback processes), cyclic graphs can also be analyzed, especially under equilibrium conditions [62]. Importantly, these causal graphs need to take into account both observed and unknown common factors (typically, these unknown factors can be and are excluded from the graph if they are all mutually exclusive [15]). In some situations, the potential confounding effects of unknown factors (context) can be eliminated using standard causal-inference techniques (e.g., using the so-called front-door and back-door criteria, or using latent variables [15]). Note that latent variables are typically used in structural equation modeling assuming linearity for all variables [17, 63]. However, when these unknown common factors cannot be eliminated or linearity cannot be assumed or validated, we propose to approximate these factors by deriving them from theoretical models (as explained in the previous section). Specifically, these unknown factors can be characterized by P_**U**_, an expected value, or can be transformed into binary variables using heuristic rules [52, 54, 57]. We provide a case study in the following section.

The translation from observational distributions to interventional distributions is rooted on *do*-calculus [15], which are the rules for moving from observations to interventions using the causal graph. That is, causal inference moves (whenever identifiable) from the probabilistic association P(y|x) to the probabilistic causal association P(y|do(x)), where *y* is the value of the potential effect *Y* and *x* is the value taken after the intervention on the inferred cause *X*. The nomenclature do(*x*) implies that we are not just merely observing *X* to take the value of *x*, but we need to make it have it (e.g., removing a species from a community). This action is then represented in a modified causal graph by eliminating all the incoming edges (causes) from an intervened variable (since its value is no longer dependent on mechanisms, but on a given action). It is typically assumed that mechanisms P(y|do(x)) are independent from each other, invariant, and follow the arrow of time (i.e., causes before effects), allowing to apply probabilistic Markov properties (i.e., each variable is independent from its non-causal variables–known as ancestors–given its causes—known as parents [15]).

Given a directed acyclic causal graph *G* and disjoint variables *X*, *Y*, *Z* and *W* (these variables can also be empty sets), do-calculus involves three rules to move from observational to interventional distributions (see Figure 1) [15]: (1) Insertion/deletion of observations: *P*(*y*|*do*(*x*), *z*, *w*) = *P*(*y*|*do*(*x*), *w*) if 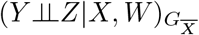, where 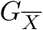 is graph *G* after the removal of all the incoming edges to *X*. This rule establishes the conditions under which it is possible to remove conditional variables from the analysis. (2) Action/observation exchange: *P*(*y*|*do*(*x*), *do*(*z*), *w*) = *P*(*y*|*do*(*x*), *z*, *w*) if 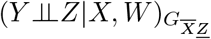, where 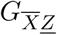 is graph 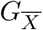 after the removal of all the outgoing edges from *Z*. This rule establishes the conditions under which it is possible to replace additional actions (acting as confounders) with observational data. (3) Insertion/deletion of actions: *P*(*y*|*do*(*x*), *do*(*z*), *w*) = *P*(*y*|*do*(*x*), *w*) if 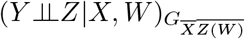, where *Z*(*W*) is the set of Z-variables that are not ancestors of any *W*-variable in 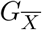. This rule establishes the conditions under which it is possible to remove additional actions (acting as confounders) from the analysis. Note that while path analysis [17] can be used instead of do-calculus, only the latter is a nonparametric framework that can be used with any sort of data without making any assumptions.

**Figure 1:**
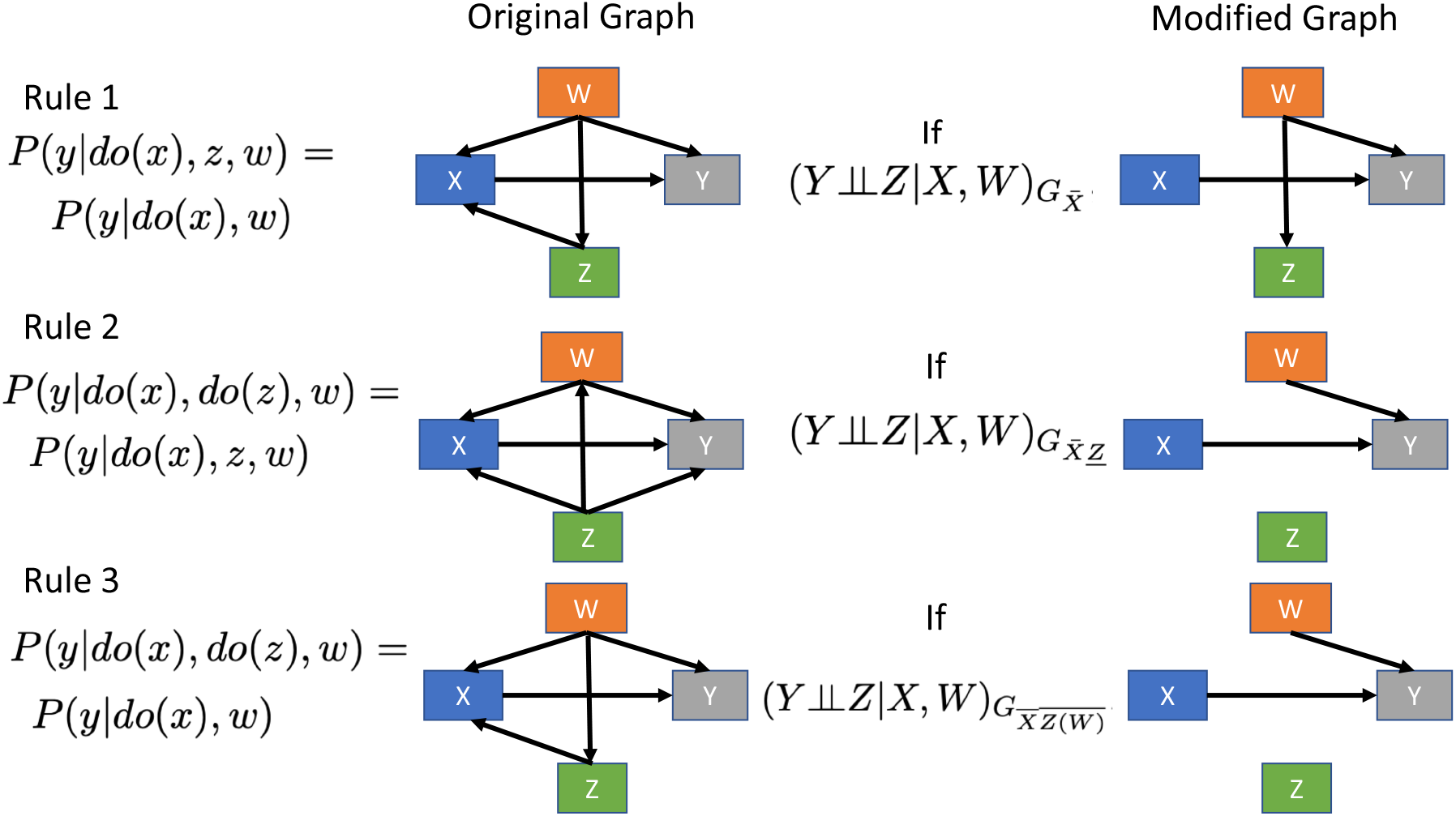
do-calculus. The translation from interventional *P*(*do*(*x*)) to observational *P*(*x*) distributions can be achieved following the rules of do-calculus [15]. The figure depicts the three do-calculus rules on a graph *G* with disjoint variables *X*, *Y*, *Z* and *W* (see main text). Rule 1 is used for insertion/deletion of observations. Rule 2 is used for action/observation exchange. Rule 3 is used for insertion/deletion of actions. Here, 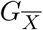 is graph *G* after the removal of all the incoming edges to *X*, 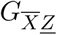 is graph 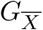 after the removal of all the outgoing edges from *Z*, and *Z*(*W*) is the set of Z-variables that are not ancestors of any *W*-variable in 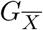. Note that 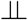 and | denote independence and conditional on, respectively. The graphs in the left column vary for illustration purposes of each rule.

## Case study

We illustrate some of the concepts above using the following example. Figure 2 depicts a hypothetical, directed, acyclic, causal graph to study the within-season pollinator abundance dynamics of a pollinator community [30, 64]. Specifically, in the example, we study how the relative abundance of flowering plants at a given time *t* (noted as *A* and measured as the ratio between the number of plant species and pollinator species at time *t*) affects the rate of change of the pollinator community at time *t* + 1 (noted as *B* and measured as the absolute difference in the pollinator community between time *t* + 1 and *t*, and divided by the observation at time *t*, providing a detrended measure). In addition, the causal graph (Fig. 2) assumes that temperature affects both *A* and *B* (written as *C* and measured as the mean temperature at time *t*). Note that *C* also works as a trend factor. Finally, we also assume that unknown factors *D* (the context) act as confounding effects of *A* and *B*. Following the concepts expressed in the previous section, we propose (see below for details) to quantify the unknown factors *D* using synthetic data derived from the linear LV model (i.e., *P*_**U**⊆**r**_) leading to the presence of the observed pollinator community at time *t* (i.e., 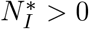). Integrating observational and synthetic data, the graph in Fig. 2 is complete and informs us about the variables that need to be blocked (controlled for) using do-calculus in order to infer the cause-effect relationships between observed variables. Note that it is assumed that each of these variables is random in the sense that they are all affected by mutually exclusive independent noise, allowing us to omit this other type of variables from the causal graph [15].

**Figure 2:**
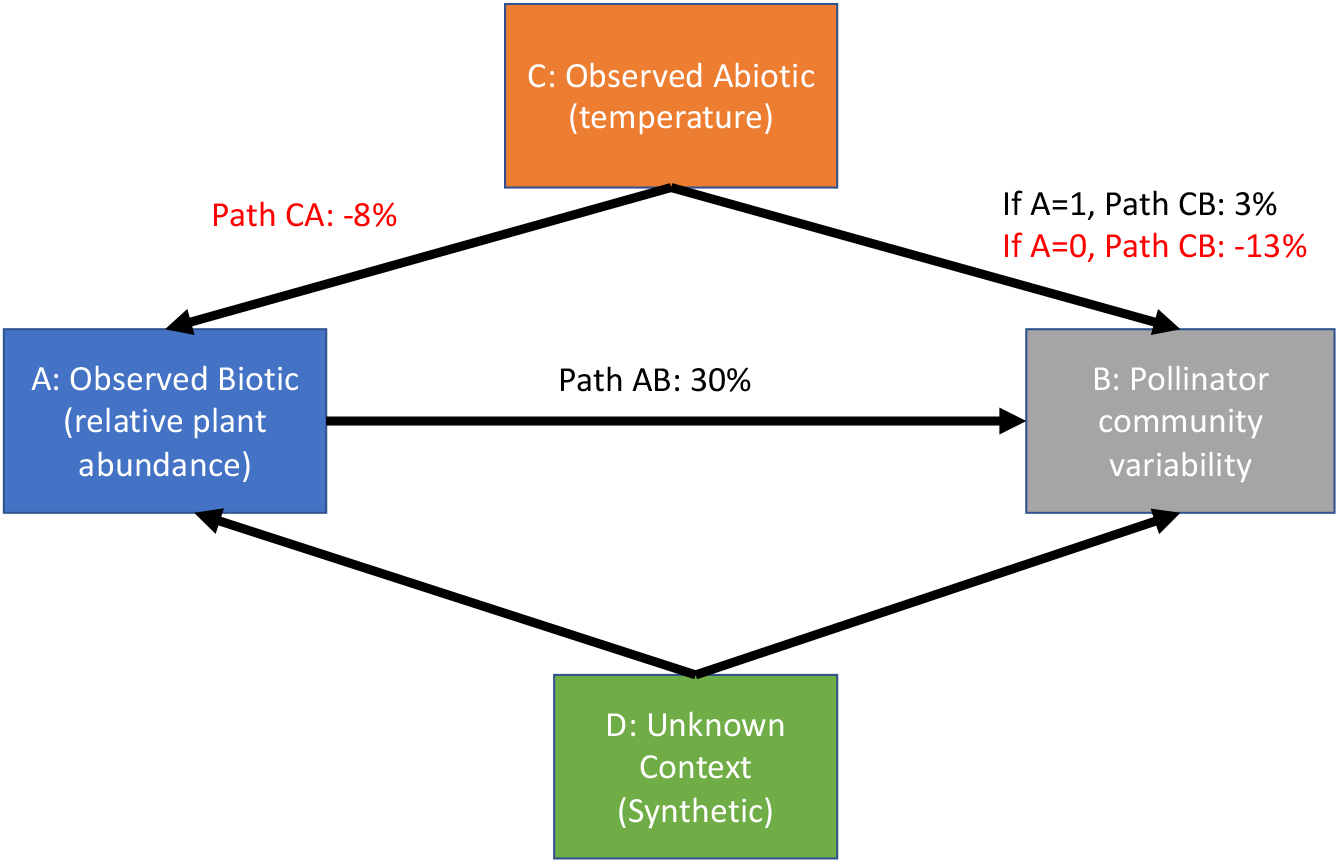
Illustrative example of cause-effect relationships of a phenological process in a pollinator community. However, this effect needs to be separated from potential confounders. The figure depicts a directed acycylic causal graph, where each box (node) corresponds to a random variable, and each edge corresponds to a direct causal effect. We consider that each causal relationship is autonomous and independent from the others. Each node is a random variable since it is also assumed that mutually exclusive random noise affects each node. Following do-calculus rules (see text and Fig. 1), for three paths, we show the estimated change in probability of observing a high value (above the median of the population) given a high value of its direct cause (see text). The variables in this graph should not be always equated to the variables in Figure 1. For example, variable C can be equivalent to variable *X* or *Z* in Figure 1 depending on the rule applied.

To put numbers to this example, we use publicly available data recording species interactions between pollinators and flowering plants on a daily basis (whenever weather allowed) in a high-arctic site during the springs of 1996 and 1997 [30, 64]. These data allow us to directly measure variables *A*, *B*, and *C* above for a given observed day *t*. To measure the theoretical context (*D*) for each day *t*, we first inferred the daily interaction matrices **A_t_** and then measure the fraction of conditions compatible with the persistence of all observed pollinators *ω*(**A_t_**). To infer **A_t_**, we use a niche-based inference [58, 65], which is one of the simplest methods yet well ecologically motivated. Specifically, we use the monopartite projection **M_t_** = **B_t_**^*T*^**B_t_**, where **B_t_** is the binary matrix for day *t* formed by the observed pollinators as columns and observed plants as rows. This binary matrix has entries *B_ki_* = 1 if the pollinator *i* is observed interacting with plant *k*, otherwise *B_ki_* = 0. In turn, the off-diagonal entries of **M_t_** correspond to the number of plant resources shared between two pollinator species. The higher the resource overlap between pollinators *i* and *j* (i.e., the value of *M_ij_*), the higher their level of competition. By normalizing the entries of **M_t_** by the sum of their column 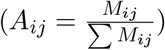, we infer a pollinator competition matrix **A_t_** for each time *t*.

To infer *ω*(**A_t_**) [30], we calculate the fraction of intrinsic growth rates (**U** ⊆ **r**) leading to the daily set of competing pollinators according to a (tractable and scalable) linear LV model. Specifically, we calculate this as:

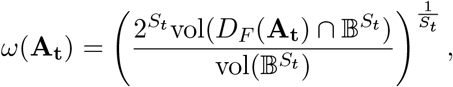

where 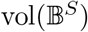 is the volume of the normalized *S_t_*-dimensional parameter space of intrinsic growth rates (**r**) at day *t*, 2*^S_t_^* normalizes the parameter space to the positive orthant (because for simplification we are summarizing the pollinator community as a competition system, all intrinsic growth rates are restricted to positive values), and 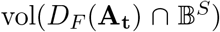 corresponds to the volume of the intersection of the the parameter space with the feasibility domain: 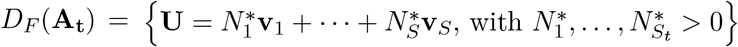, where **v**_*i*_ is the *i*th column vector of the in teraction matrix **A_t_** [54]. Thus, *ω*(**A**_*t*_) ∈ [0, 1] is a probabilistic measure, which can be efficiently computed and compared across dimensions [30, 54].

Similar to path analysis in structural equation modeling [17, 63], to apply probabilistic causal inference with continuous data, it can be possible to use linear regressions (or Pearson correlations) if it is assumed that the effects are linear, monotonic, and noise is Gaussian. Spearman rank correlations can be used if at least monotonicity is achieved. Instead, nonparametric tools can be used whether or not these assumptions above are fulfilled. While nonparametric tools provide generality and should be preferred, their application to continuous data can be rather challenging. Thus, whenever possible, the data can be discretized [15]. Here, for illustration purposes, we transform all our variables into binary values, using the median of each variable (per year) as the cut-off value: values higher that the median are considered one, otherwise zero. While this may be perceived as a disadvantageous simplification, it actually allows us to efficiently work on a general nonparametric framework (i.e., using probability distributions).

We test the causal graph shown in Figure 2. Here, the only testable d-separation (conditional or unconditional) is between temperature (*C*) and context (*D*). That is, there is no direct path between these two variables, and their path gets naturally blocked (no need to condition on anything) by *A* and *B*, which act as colliders. This d-separation can be tested by the unconditional independence as *P*(*d*|*c*) = *P*(*d*). Using a *G*^2^-test (*χ*^2^-test can also be used for binary data or permutation tests [15, 17]), we found no statistical relationship between *C* and *D* (*p* = 0.39, lower values indicate dependence). Note that if the hypothesis would not have been supported by d-separation, a new causal graph must be drawn and tested. Below, we compute the effects of temperature on the relative abundance of flowering plants (Path CA), the effect of temperature on community variability (Path CB), and the effect of relative abundance of flowering plants on community variability (Path AB).

The interventional distribution (probability of cause) of Path CA is written as *P*(*a*|*do*(*c*)). This causal relationship can be inferred using observational distributions following rule 2 of do-calculus. That is, we can write *P*(*a*|*do*(*c*)) = *P*(*a*|*c*) by setting *Y* = *A*, *Z* = *C*, and 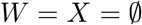 in Figure 1. Because we are using binary variables, the average causal effect [15] of *c* on *a* (i.e., *ACE_CA_*) is given by 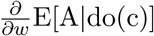 and can be written as *ACE_CA_* = *P*(*a* = 1|*c* = 1) – *P*(*a* = 1|*c* = 0). We found that *ACE_CA_* = −0.08, meaning that if temperature is high (i.e., above the population median) there is a decrease in probability of 8% that the relative plant abundance will be high (i.e., above its population median). However, using a *G*^2^ test, we found that this effect is not largely different (*p* = 0.56) from what would be expected by chance alone given the data. In turn, the interventional distribution of Path CB can be calculated as *P*(*b*|*do*(*c*), *do*(*a*)). Note that Path CB is mediated by *A*, which needs to be controlled for. However, conditioning (i.e., *P*(*b*|*do*(*c*), *x*)) opens the collider between *C* and *D*, creating a spurious association between *C* and *B*. To eliminate this noise, it is then necessary to intervene on *A* (i.e., *do*(*a*)). Using marginalization and the Markov property, we can write *P*(*b*|*do*(*c*), *do*(*a*)) = ∑_*d*_ *P*(*b*, *d*|*do*(*c*), *do*(*a*)) = ∑_*d*_ *P*(*b*|*do*(*c*), *do*(*a*), *d*)*P*(*d*). Following rule 2 twice (setting first *Z* = *A*, *Y* = *B*, *X* = *C*, and *W* = *D*; and second *Z* = *C*, *Y* = *B*, 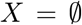, and *W* = {*A*, *D*} in Fig. 1), we can write ∑_*d*_ *P*(*b*|*c*, *a*, *d*)*P*(*d*). In this case, we can perform two separated analyses: one for *a* = 1 and the other for *a* = 0. We found that for *a* = 1, *ACE_CB_* = 0.03 (*G*^2^ test: *p* = 0.43). While for *a* = 0, *ACE_CB_* = −0.13 (*G*^2^ test: *p* = 0.008). This implies that under high flower abundance, temperature has almost no effect on pollinator variability. Instead, under low flower abundance, if temperature is high (i.e., above the population median), there is a decrease in probability of 13% that the variability of the pollinator community will be also high (i.e., above its population median).

Finally, following the methodologies above, we calculate the effect of relative plant abundance on community variability (Path AB) as *P*(*b*|*do*(*a*)) = ∑*_cd_ P*(*b*|*a*, *c*, *d*)*P*(*c*, *d*). We found that *ACE_AB_* = 0.30 (*G*^2^ test: *p* = 0.06), meaning that if relative plant abundance is high (i.e., above the population median) there is an increase in the probability of 30% that the community variability will be high (i.e., above its population median). It is worth mentioning that if we do not take into account the context (*D*), the causal effect of *A* (relative flower abundance) on *B* (pollinator community variability) can be overestimated *ACE_AB_* = 0.86 (*G*^2^ test: *p* = 0.003), leading to potential prediction errors of interventions. It is also important to mention that a linear multivariate regression of *B* on all the other three variables (using normalized data instead of binary) produce qualitatively similar results as the ones reported above. While this equivalence between nonparametric and parametric methods is not expected to be always true [15], working under a causal hypothesis (as we have done here) can establish a more informative regression analysis that can then be translated into causal analysis under the assumption of linearity.

This example is not intended to demonstrate a general effect and serves only for illustration purposes. For example, we try to explain a fairly simple community metric such as changes in overall relative abundance. Furthermore, many more variables can be explicitly taken into account (instead of being summarized in the unknown confounding factors), such as abundance of pathogens, herbivores, chemical compounds, humidity, etc, and it is important to identify the main players in line with the hypothesized causal graphs. Moreover, it is important to note that the theoretical model has also sensible assumptions, such as that resource overlap among pollinators is a good proxy of competition. We hope future work can build on this to establish causal knowledge at the pollinator community-level.

## Conclusions

It has long been recognized that causation does not always coincides with correlation. This premise has been extensively applied when studying the behavior (i.e., variables) of complex natural systems, where multiple factors can be responsible for the patterns observed in nature. This has not been an exception when investigating pollinator communities. As a consequence, the majority of work has carefully stated correlations, which respond to what do we see in nature. However, in the face of rapid environmental change, we need to take bolder research programs and answer the questions of why and when the behavior of pollinator communities is affected. These goals can be achieved by conducting experimental studies. Nevertheless, manipulating all factors related to the behavior of entire pollination communities can be unrealistic. Instead, these goals can be achieved by using causal-inference techniques. Yet, very often these techniques cannot be applied due to the nature of the causal story and the unknown/unmeasured factors acting as confounders. While not exhaustive, here we have provided a brief overview of how to apply probabilistic causal inference from the integration of observational and synthetic data. We propose that synthetic data can be used as a proxy for unknown confounding factors by deriving them from theoretical models that attain the desired properties of tractability (provide a systematic link between model parameters and solutions) and scalability (can be applied across dimensions). At the very least, we hope this overview can illustrate that a causal probabilistic analysis can allow us to speak the causal language in pollination studies that for long has been prevented by the dominance of multivariate regressions and meta-analyses without causal hypotheses [32].

## Acknowledgments

Funding to SS was provided by NSF grant No. DEB-2024349. IB and OG acknowledes funding from the Simplex project (PRPCGL2017-92436-EXP). OG acknowledges financial support provided by the Spanish Ministry of Economy and Competitiveness (MINECO) and by the European Social Fund through the Ramón y Cajal Program (RYC-2017-23666). RPR acknowledges funding from the Swiss National Science Foundation, grant no. 31003A_182386. PZ acknowledges the Swiss National Science Foundation Spark scheme, under grant number CRSK-3_196506.

## Author contributions

SS designed and performed the study. All authors contributed with ideas and wrote the manuscript.

## Data accessibility

The data and R code supporting the results can be found at https://github.com/MITEcology/Saavedra_etal_causal_example.

